# Notch regulated long non-coding RNA TUG1 regulates smooth muscle cell differentiation in aortic aneurysm

**DOI:** 10.1101/2023.04.21.537806

**Authors:** Ravi Abishek Bharadhwaj, Regalla Kumarswamy

## Abstract

Abdominal Aortic Aneurysms (AAAs) are asymptomatic vascular diseases with life threatening outcomes. Smooth-muscle cell (SMC) dysfunction plays an important role in AAA development. The contributions of non-coding genome, specifically the role of long non-coding RNAs (lncRNAs) in SMC dysfunction are relatively unexplored. We investigated the role of lncRNA TUG1 in the pathology of AAA. TUG1 was identified through lncRNA profiling in Angiotensin-II (Ang-II) treated SMCs. TUG1 was upregulated in Ang-II treated SMCs *in vitro* and its expression increased with progression of aneurysm in mouse model of Ang-II induced AAA. Ang-II induced TUG1 was blunted by inhibition of Notch signaling and TUG1 is demonstrated to be a transcriptional target of Notch. AAA tissues exhibited inversely correlated expression of TUG1 and SMC contractile markers. TUG1 knock-down via siRNA/shRNA increased SMC differentiation. ChIP, DNA-RNA IP, and RNA-IP experiments demonstrated that TUG1 interacts with transcriptional repressor KLF4 and aides in its recruitment to Myocardin promoter, thereby repressing SMC differentiation. In summary, we show a novel role for lncRNA TUG1 in Ang-II induced AAA wherein it modulates SMC differentiation via KLF4-Myocardin axis.

## Introduction

An aortic aneurysm is characterized by permanent localized dilatation of aorta to more than 1.5 times of normal aortic diameter. No medical interventions have been validated to attenuate expansion or prevent rupture. Based on the location, aortic aneurysms are classified as thoracic aortic aneurysm (TAA) or abdominal aortic aneurysm (AAA) (Quintana and Taylor, 2019). Previous studies reported that prevalence of AAA ranges up to 7.2% in general population aged 60 or above (Guirguis-Blake *et al*., 2019). At the histopathological level, AAA is characterized by smooth muscle cell (SMC) dysfunction, extracellular matrix (ECM) degradation and aortic wall inflammation, all of which lead to the degeneration of vessel wall and loss of aortic integrity. SMCs in medial layer are the most abundant cells of the aorta and are known to secrete key ECM proteins that maintain aortic contraction and vasotone (Chen *et al*., 2020). SMCs, unlike other mature cells, retain their plasticity and exhibit the ability to undergo phenotypic switch during development and also during any form of vascular injury (Espinosa-Diez *et al*., 2021). The importance of these phenotypic transitions of SMCs has been well explored in evolution of atherosclerotic plaques (Allahverdian *et al*., 2018). Earlier studies have also shown that SMC phenotypic modulation is an early event of aortic aneurysm formation (Ailawadi *et al*., 2009), and SMCs from aneurismal aortas exhibit reduced contractility (Bogunovic *et al*., 2019). However, the molecular mediators of pathophysiological events in SMCs during AAA are still not well understood. Long non coding RNAs (lncRNAs) are a class of non coding RNAs that are more than 200 bp in length. They regulate several processes including cell proliferation (Ahmed *et al*., 2018), stem cell pluripotency (De Jesus *et al*., 2018) and cell differentiation (Dong *et al*., 2021). Several lncRNAs are also known to regulate SMC function in development and vascular diseases. Recently, MALAT1 was shown to mediate SMC dysfunction in TAA through interaction with HDAC9-BRG1 (Lino Cardenas *et al*., 2018), other examples include H19 (Li *et al*., 2018), SENCR (Bell *et al*., 2014), SMILR (Ballantyne *et al*., 2016), Lnc-Ang362 (Leung *et al*., 2013) and GAS5 (He *et al*., 2019) which are shown to influence SMC physiology. Taurine Upregulated Gene1 (TUG1), a 6.7 kb conserved lncRNA, plays important roles in diabetic nephropathy (Long *et al*., 2016) and cancers (Li *et al*., 2016). Targeting Notch-TUG1 axis was proposed as specific therapeutic approach for glioblastoma (Katsushima *et al*., 2016). In the context of vascular diseases, TUG1 is known to negatively regulate SMC function in atherosclerotic events (Chen *et al*., 2017; Li, Lin and Gao, 2018; Shi, Zhu and Fan, 2021). However, the role of TUG1 in AAA is not known.

In this study, we investigated the role of TUG1 in Angiotensin-II (Ang-II) induced AAA in Apolipoprotein E deficient (ApoE^-/-^) mice. Using *in vitro* and *in vivo* models, we show that Ang-II induced TUG1 is involved in SMC dysfunction. Using complementing immunoprecipitation techniques, we established a novel link between TUG1 and KLF4-Myocardin axis, which is crucial in maintaining SMC phenotype.

## Results and Discussion

### TUG1 is an Angiotensin-II induced long non coding RNA in smooth muscle cells

To identify differentially expressed long non coding RNAs (lncRNAs) in smooth muscle cells (SMCs) that are relevant to abdominal aortic aneurysm, we treated mouse aortic SMCs with Angiotensin-II and performed lncRNA profiling for 84 well characterized lncRNAs that are previously implicated in human diseases (Qiagen, Mouse lncFinder RT2 lncRNA PCR Array, Cat no: 330721 LAMM-001Z). Out of 84 lncRNAs, 45 lncRNAs were expressed in SMCs, out of which 27 lncRNAs were deregulated with a fold change of 1.5 or more (Fig S1A). Among these TUG1 and GAS5 are vascular enriched and conserved in mouse, rat and humans (Fig S1B, Supplementary Table 1). The role of GAS5 has already been explored in AAA (He *et al*., 2019). Deregulation of TUG1 in multiple aortic SMCs upon Angiotensin-II treatment was further verified in independent qPCR assays (Fig 1A). Higher expression of TUG1 was found in SMC rich tissues such as aorta, brain and lungs (Fig S1C).

**Figure 1:**
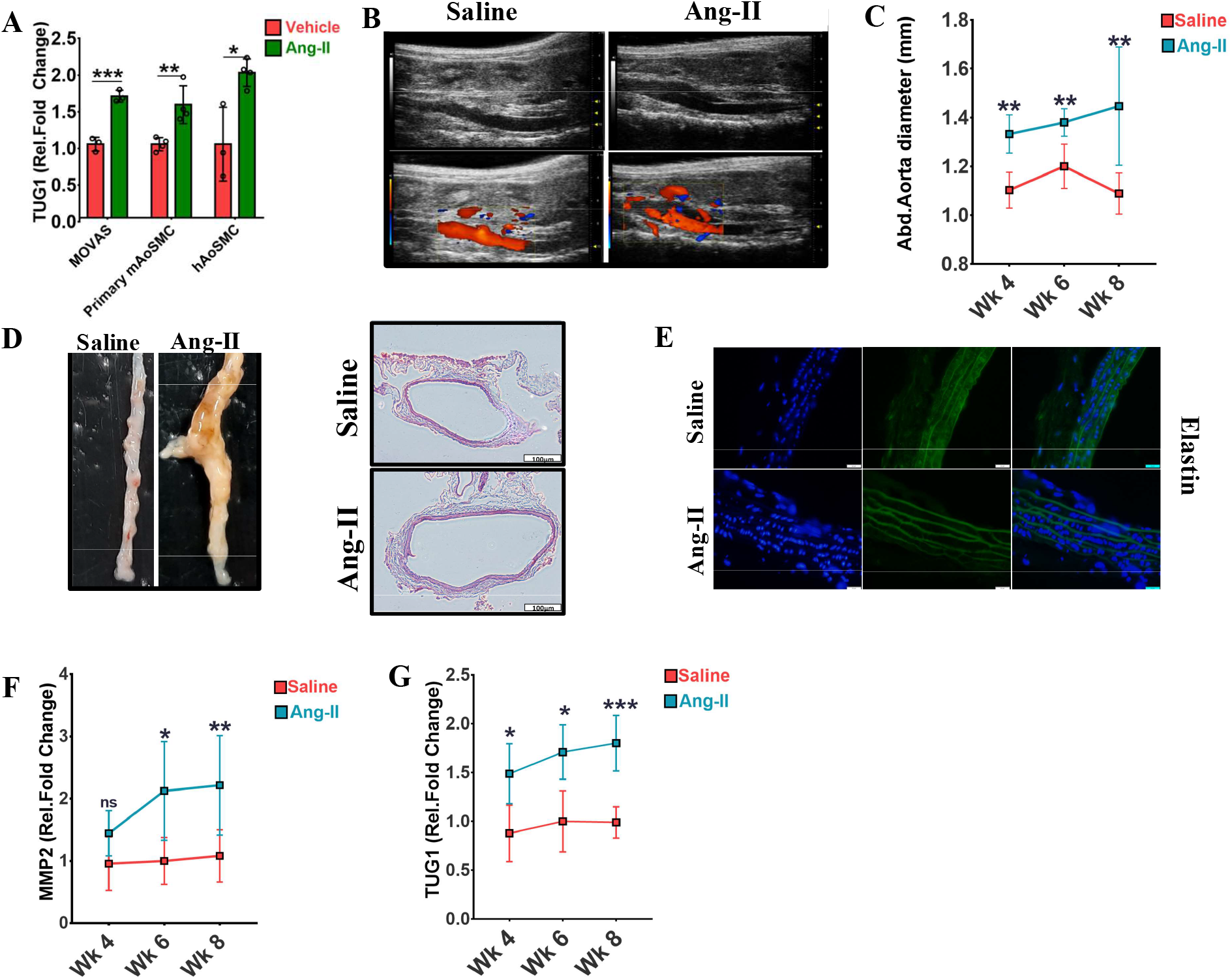
TUG1 is an Ang-II induced long non coding RNA involved in abdominal aortic aneurysm. (A) TUG1 expression was profiled in mouse aortic smooth muscle cells (MOVAS), primary mouse aortic smooth muscle cells (Primary mAoSMC) and human aortic smooth muscle cells (hAoSMCs) upon Angiotensin-II treatment (Ang-II) for 24 h, data was shown as relative fold change over control after normalization with RPLP0 and depicted as mean ± SD, n=3-4/group. (B) Representative ultrasound image of Angiotensin-II (Ang-II) induced abdominal aortic aneurysm (AAA) at 8 weeks. (C) Abdominal aorta (Abd.Aorta) diameter progression during development of AAA, data is shown as mean of Abd.Aorta diameter ± SD, n=5-7/group. (D) Representative image of dissected abdominal aortas at 8 weeks and H&E stained cross section of Abd.Aorta, scale bar 100 μm. Please note dilation of the aorta and increased lumen size in Ang-II treated animals. (E) Immunoflourescent elastin staining of Abd.Aorta at 8 weeks, scale bar: 20 μm. (F) qPCR quantification of MMP2 and TUG1 (G). Data shown as relative fold change over saline treated abdominal aortas after normalization with GAPDH/RPLP0 and depicted as mean ± SD, n=3-5/group. * Denotes p<0.05, ** denotes p<0.01, *** denotes p<0.001

### TUG1 is upregulated during Angiotensin-II induced abdominal aortic aneurysm (AAA) in mice

AAA was induced by administration of Angiotensin-II for 8 weeks in ApoE-/- mice as described in Material and Methods. Development of aneurysm was monitored using small animal ultrasound imaging system (Fig 1B). Abdominal aortas of Angiotensin-II treated mice displayed increased abdominal aortic diameter from week 4 onwards (Fig 1C). Histological analysis of aneurismal tissues revealed increased lumen diameter (Fig 1D), reduced elastin content (Fig 1E) and break in elastin fibers compared to saline aortas (Fig S2A). MMP2, which is an indicator of aneurysm development, was upregulated in abdominal aortas of Angiotensin-II treated mice (Fig 1F). In line with cultured SMCs, TUG1 was upregulated in abdominal aortas in mice during Angiotensin-II induced aneurysm (Fig 1G).

### TUG1 expression is regulated by Notch signaling

Notch signaling is previously known to mediate Angiotensin-II induced vascular remodeling (Ozasa *et al*., 2013) and inhibition of Notch attenuated Angiotensin-II induced AAA (Cheng *et al*., 2014; Sharma *et al*., 2019). To understand if Notch has any role in TUG1 upregulation, mouse aortic SMCs were treated with Angiotensin-II alone and in combination with Notch pharmacological inhibitor N-[N-(3,5-Difluorophenacetyl)-L-alanyl]-S-phenylglycine t-butyl ester (DAPT). As shown in Fig 2A upregulation of TUG1 after Angiotensin-II treatment was blunted by DAPT. Hey1, a known transcriptional target of Notch signaling served as a positive control for Notch activation. Similarly, overexpression of Notch1 intra-cellular domain (Notch1-ICD) increased TUG1 expression (Fig 2B) while inhibition of Notch1 with siRNA decreased TUG1 expression (Fig 2C). Moreover, Angiotensin-II mediated increase in TUG1 in aneurismal aortas was also blunted in animals co-treated with DAPT (Fig 2D). As Notch does not have a DNA binding domain, Notch specific gene activation is mediated through Notch binding with RBPJK which in turn binds to DNA (Wang *et al*., 2012; Castel *et al*., 2013). We examined TUG1 promoter for presence of RBPJK binding sites and found potential binding (Fig 2E). Luciferase reporter assays with TUG1 promoter revealed that activation of Notch signaling by over expressing Notch1-ICD induced luciferase activity (Fig 2F) while Notch inhibition with DAPT had an opposite effect (Fig 2G). Chromatin immunoprecipitation also confirmed Notch1-ICD enrichment on TUG1 promoter (Fig 2H).

**Figure 2:**
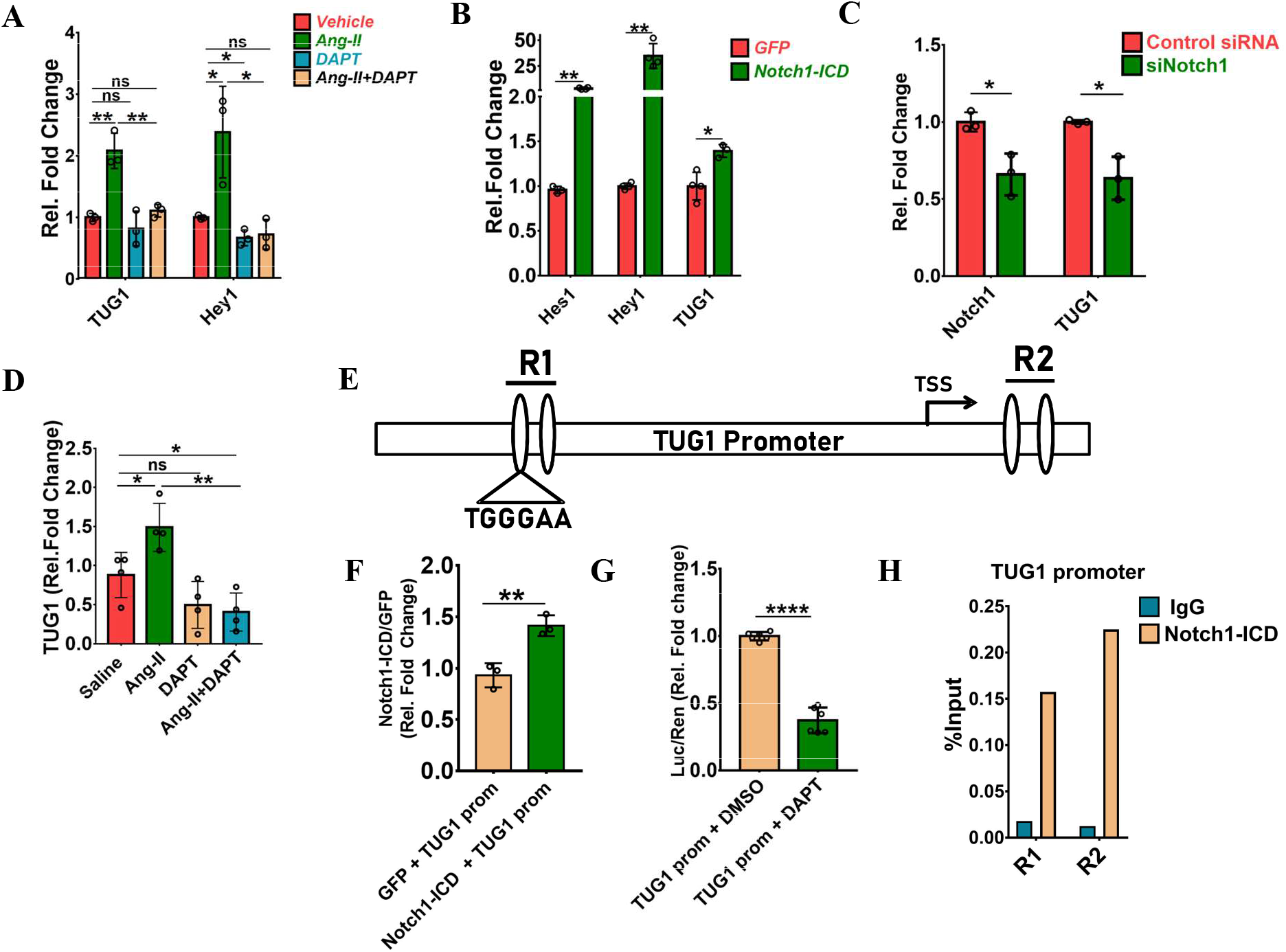
TUG1 is an Ang-II induced Notch regulated lncRNA. (A) TUG1 expression after 24 h in mouse aortic smooth muscle cells (MOVAS), treated with Angiotensin-II (Ang-II) (1 μM) alone or in combination with Notch signaling inhibitor N-[N-(3,5-Difluorophenacetyl)-L-alanyl]-S-phenylglycine t-butyl ester (DAPT) (10 μM). Hey1 was used a positive control of Notch signaling, data is shown as relative fold change over control, after normalization with RPLP0 and depicted as mean ± SD, n=3/group. (B) TUG1 expression after 24 h in MOVAS overexpressing Notch1-Intracellular domain (Notch1-ICD), Hey1 and Hes1 were used a positive controls of Notch signaling, data was shown as relative fold change over control after normalization with RPLP0 and depicted as mean ± SD, n=3-4 /per group. (C) TUG1 expression after 24 h in MOVAS upon Notch1 KD with siRNA (50 nM), data is shown as relative fold change over control siRNA after normalization with RPLP0 and depicted as mean ± SD, n=3/group. (D) TUG1 expression in aortas treated with Angiotensin-II alone or in combination with Notch inhibitor DAPT (10mg/kg/day, 3 days/ week). Data was shown as relative fold change over saline treated abdominal aortas after normalization to GAPDH and depicted as mean ± SD, n=4/group. (E) Representative image of TUG1 promoter with Notch-RBPJK DNA elements R1 and R2 annotated. (F) TUG1 promoter luciferase reporter assay performed after 24 h in MOVAS overexpressing Notch1-ICD, data is shown as relative fold change of Notch1-ICD rel.GFP and depicted as mean ± SD of Luc/Ren values, n=3/group. (G) TUG1 promoter luciferase reporter assay was performed after 24 h in MOVAS treated with Notch signaling inhibitor DAPT (10 μM), data is shown as relative fold change over control and depicted as mean ± SD of Luc/Ren values, n=6/group. (H) Chromatin immunoprecipitation of TUG1 promoter was performed using Notch1-ICD antibody in ectopic Notch1-ICD overexpressing cells, data is shown as enrichment % input. * Denotes p<0.05, ** denotes p<0.01, *** denotes p<0.001, **** denotes p<0.0001.

### TUG1 regulates SMC differentiation

In response to injury, SMCs undergo profound phenotypic change to a de-differentiated state characterized by repression of smooth muscle marker genes and reduced contractility (Allahverdian *et al*., 2018; Frismantiene *et al*., 2018; Qian *et al*., 2022). SMC phenotypic modulation is an early event of AA formation (Ailawadi *et al*., 2009), and SMCs from AAAs exhibit reduced contractility (Bogunovic *et al*., 2019). As shown in Fig 3A, TUG1 is upregulated in animals upon Angiotensin-II treatment while smooth muscle markers SM22-α and α-SMA are repressed. Immunostaining of AAA tissues also revealed reduced expression of SMC marker genes SM22α, CNN1 and α-SMA (Fig 3B). Silencing of TUG1 resulted in upregulation of differentiation markers Myocardin, Smoothelin, SM22α and PTEN (Fig 3C). This also led to increased contractility as evidenced from collagen gel contraction assay in TUG1 KD cells (Fig 3D).

**Figure 3:**
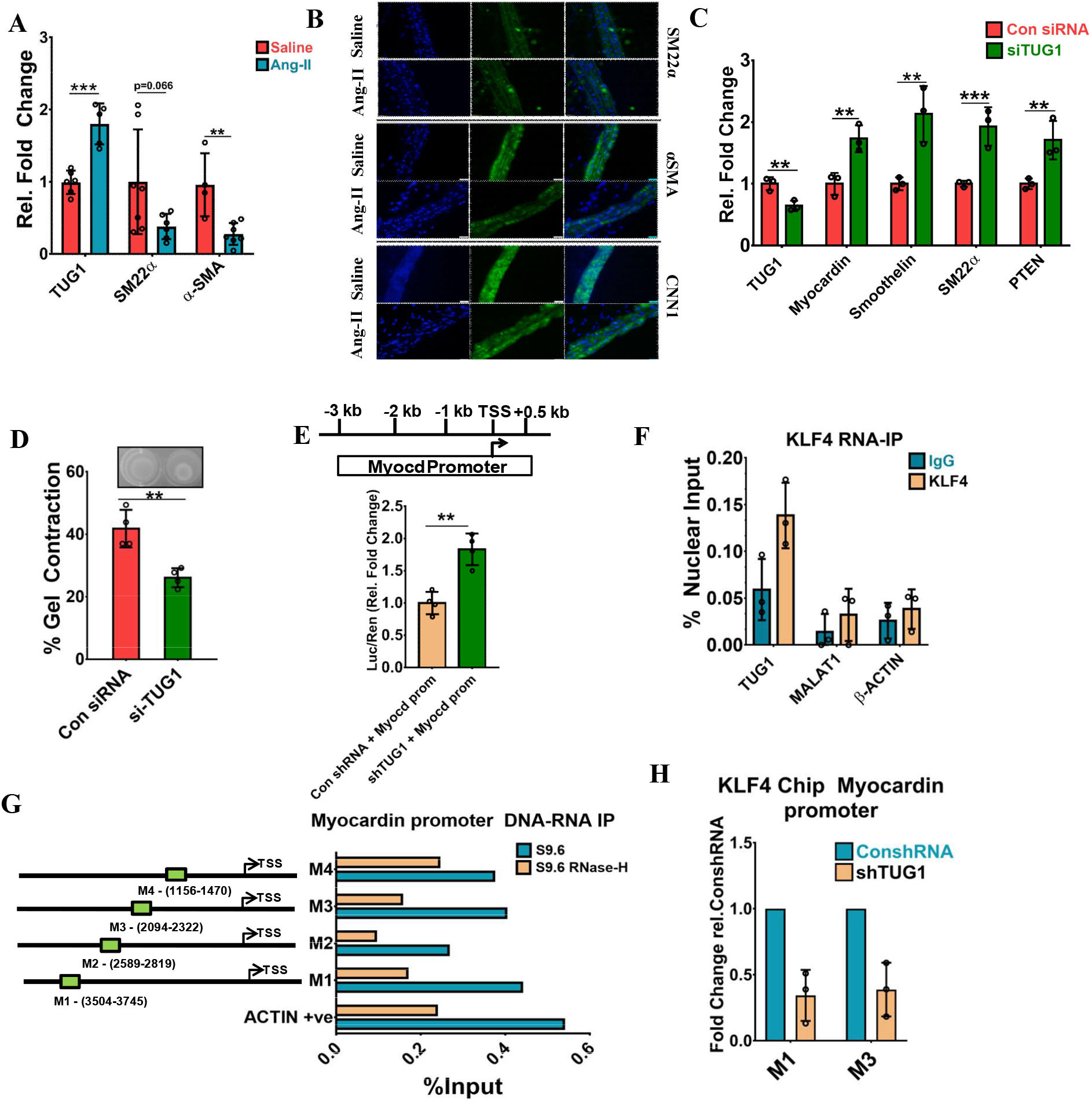
TUG1 regulates SMC differentiation through Myocardin via interaction with KLF4. (A) qPCR expression of TUG1 and SMC differentiation markers α-SMA and SM22α in AAA tissues after 8 weeks of Angiotensin-II (Ang-II) administration, data is shown as relative fold change over saline treated abdominal aortas normalized to RPLP0/GAPDH and depicted as mean ± SD, n=4-5 /group. (B) Immunofluorescence staining of abdominal aortas of aneurismal and saline treated abdominal aortas stained for SMC differentiation markers SM22α, α-SMA and CNN1, images depicted were acquired at 40X magnification, Scale bar: 20μm. (C) qPCR expression of SMC differentiation markers 24 h after siRNA (100nM) mediated TUG1 silencing in mouse aortic SMCs (MOVAS), data is shown as relative fold change over negative control siRNA normalized to RPLP0 and depicted as mean ± SD, n=3/group. (D) Collagen gel contraction assay in si-TUG1 KD in MOVAS, data is shown as % contraction of gel relative to decellularized gel and depicted as mean ± SD, n=4 /group. (E) Mouse Myocardin promoter used in luciferase assay. Myocardin promoter (Myocd prom) luciferase reporter assay was performed after 48 h in TUG1 KD mouse aortic SMCs, data was shown as relative fold change over negative control shRNA Luc/Ren values and depicted as mean ± SD, n=4/group. (F) TUG1 RNA immunoprecipitation was performed with KLF4 antibody in MOVAS, data is shown as % of Nuclear Input. Nuclear lncRNA MALAT1 and β-Actin were used as negative controls and depicted as mean ± SD of 3 independent pulldowns. (G) DNA-RNA immunoprecipitation of Myocardin promoter (representative image of sites profiled), Actin locus was used as positive control of DNA-RNA IP, data is shown as % Input. (H) Chromatin immunoprecipitation of Myocardin promoter was performed with KLF4 antibody in TUG1 KD cells, data is shown as % relative enrichment over negative control shRNA after normalization with IgG, n=3 independent pulldowns. * Denotes p<0.05, ** denotes p<0.01, *** denotes p<0.001

shRNA mediated silencing of TUG1 (Fig S3A) also increased promoter activity of Myocardin in mouse aortic SMCs (Fig 3E). Myocardin is a potent SMC phenotypic modulator (Pipes, Creemers and Olson, 2006; Long *et al*., 2007, 2008; Parmacek, 2008; Hoofnagle *et al*., 2011) whose expression is regulated by transcriptional repressor KLF4 (Liu *et al*., 2005). In silico analysis using catRAPID tool (Bellucci *et al*., 2011) revealed that TUG1 has high propensity to bind with KLF4 (Fig S3B). RNA-immunoprecipitation confirmed TUG1-KLF4 interaction (Fig 3F). Further, cell fractionation studies also confirmed TUG1 to be majorly chromatin enriched (Fig S3C). To see if TUG1-KLF4 interaction has any implication for regulation of Myocardin expression, we performed RNA-DNA immunoprecipitation in mouse aortic SMCs with anti-DNA-RNA hybrid antibody S9.6 and found Myocardin promoter harbors three RNA binding elements shown as M1, M2 and M3 (Fig 3G). Sequence of these sites revealed that sites M1 and M2 harbor core KLF4-DNA binding motif GGTG, (Liu *et al*., 2014), while M3 was found to resemble a previously described motif of TUG1 DNA binding motif (A_50_) (Long *et al*., 2016). To test if TUG1 interaction with KLF4 aides in the recruitment of the later to Myocardin promoter, we performed pulldown of Myocardin promoter with KLF4 in TUG1 KD cells and found reduced enrichment at M1 and M3 (Fig 3H). Taken together, these data indicate that TUG1 interacts with transcriptional repressor KLF4 and facilitates its recruitment to Myocardin promoter leading to repression of smooth muscle marker genes and reduced contractility, which are hallmarks of SMC dysfunction (Fig 4).

**Figure 4:**
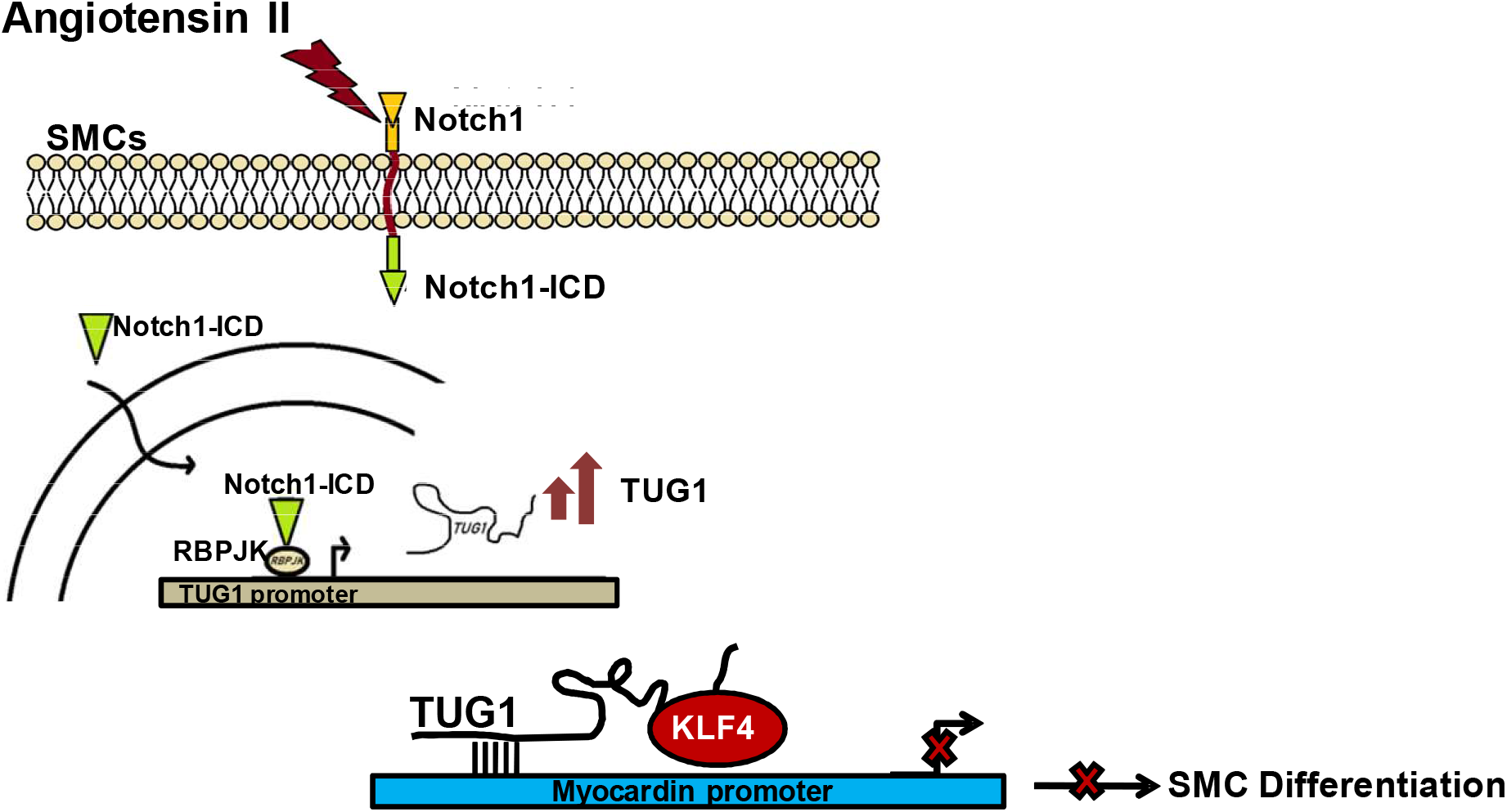
Working model. Angiotensin-II induces TUG1 through Notch signaling. TUG1 interacts with transcriptional repressor KLF4 and helps in its recruitment to Myocardin promoter resulting suppression of SMC differentiation and contractile genes.

## Material & Methods

### Mouse model of Abdominal Aortic Aneurysm (AAA)

All animal experiments were conducted as per institutional animal ethics committee guidelines and ARRIVE guidelines. AAA was induced as previously described (Daugherty, Manning and Cassis, 2000) with some modifications. 12-14 wk old Apolipoprotein E deficient (ApoE^-/-^) male mice were administered with Angiotensin-II, (Sigma, Cat No A9525) (1 mg/kg/day) or saline for 56 days. For Notch inhibition, animals were injected with N-[N-(3,5-Difluorophenacetyl)-L-alanyl]-S-phenylglycine t-butyl ester (DAPT) (Sigma, Cat no. D5942) (10 mg/kg/day, 3 times a week). The development of aneurysm was monitored using Vevo 2100 small animal echocardiography system (FujiFilm Visual sonics). For measurement of aneurismal dilation, aortic diameters at three different segments of the abdominal aorta in diastole were analyzed using Vevo lab software (FujiFilm visual sonics). After confirming the development of AAA, aortas were harvested and abdominal aortas (until iliac bifurcation) were used for qPCR studies or histology. Antibody and their dilutions used in immunoflourescent staining are mentioned in Table 2. All microscopy images were acquired using Olympus IX73 inverted microscope (Evident scientific).

### Cell Culture treatments

Mouse aortic smooth muscle cells (ATCC CRL 2797- MOVAS) cells were grown in DMEM medium and starved in 0.2 % serum unless mentioned otherwise. All starvations were performed 24 h prior to treatment. Cells were treated with Angiotensin-II (1 μM) alone or in combination with DAPT (10 μM) or siNotch1 (Qiagen) (50 nM) or siTUG1 (Qiagen) (100 nM) as indicated in results section. Primary mouse aortic smooth muscle cells were isolated as previously described (Adhikari *et al*., 2015) and grown in DMEM. Human aortic smooth muscle cells (Promocell Cat No – C12533) were grown in M231 medium. Human and primary mouse aortic SMCs were starved in 0 % serum for 48 h prior to treatment. For Notch overexpression, Notch1 intracellular domain (Notch1-ICD) was cloned in GFP expressing vector. Transfections were carried out with Lipofectamine 3000 or RNAi Max as per manufacturer instructions.

Sequences of si-RNA/shRNA used are mentioned in Table 3. All cell cultures were tested for mycoplasma contamination prior to treatments.

### Collagen gel contraction assay

Collagen gel contraction assay was performed as per Ngo *et al*., 2006 in mouse aortic smooth muscle cells that were treated with siTUG1 for 24 h prior to embedding in Collagen gel lattices, (Collagen type I, Thermofisher, Cat No - 1048301). Images were acquired 48h later and gel contraction was quantified with Image J.

### Promoter Luciferase reporter assay

Luciferase reporter assays were performed as per manufacturer’s instructions (Dual luciferase Assay kit, Cat no - E1910, Promega). Mouse TUG1 promoter DNA sequence comprising -3000 bp and +500 bp to TSS was cloned in pGL3 basic luciferase vector. Mouse aortic smooth muscle cells were co-transfected with TUG1 promoter and Notch1-ICD over-expressing vectors to study TUG1 promoter activity upon Notch activation. For studying TUG1 promoter activity upon Notch inhibition, mouse aortic smooth muscle cells were pre-treated for 24 h with DAPT (10 μM). For Myocardin promoter assays, mouse Myocardin promoter DNA sequence comprising -3000 bp and +500 bp to TSS was cloned into pGL3 basic luciferase vector. Cells were transfected with reporter vector in control shRNA or shTUG1 cells as indicated in the results. All transfection experiments were performed with 1000 ng of respective plasmid in 24 well plate using Lipofectamine 3000. pRL-SV40 renillia luciferase vector served as normalizing control.

### Sub-Cellular Fractionation

Cytoplasmic and nuclear fractions were separated as previously described (Conrad and ørom, 2017) using density gradient centrifugation. RNA from cell fractions was isolated using Trizol reagent. Transcript enrichment was studied using Sybr-green qPCR. NEAT1 and MALAT1 were used as positive controls for nuclear fractions. Primer sequences are mentioned in Table 1.

**Table 1.**
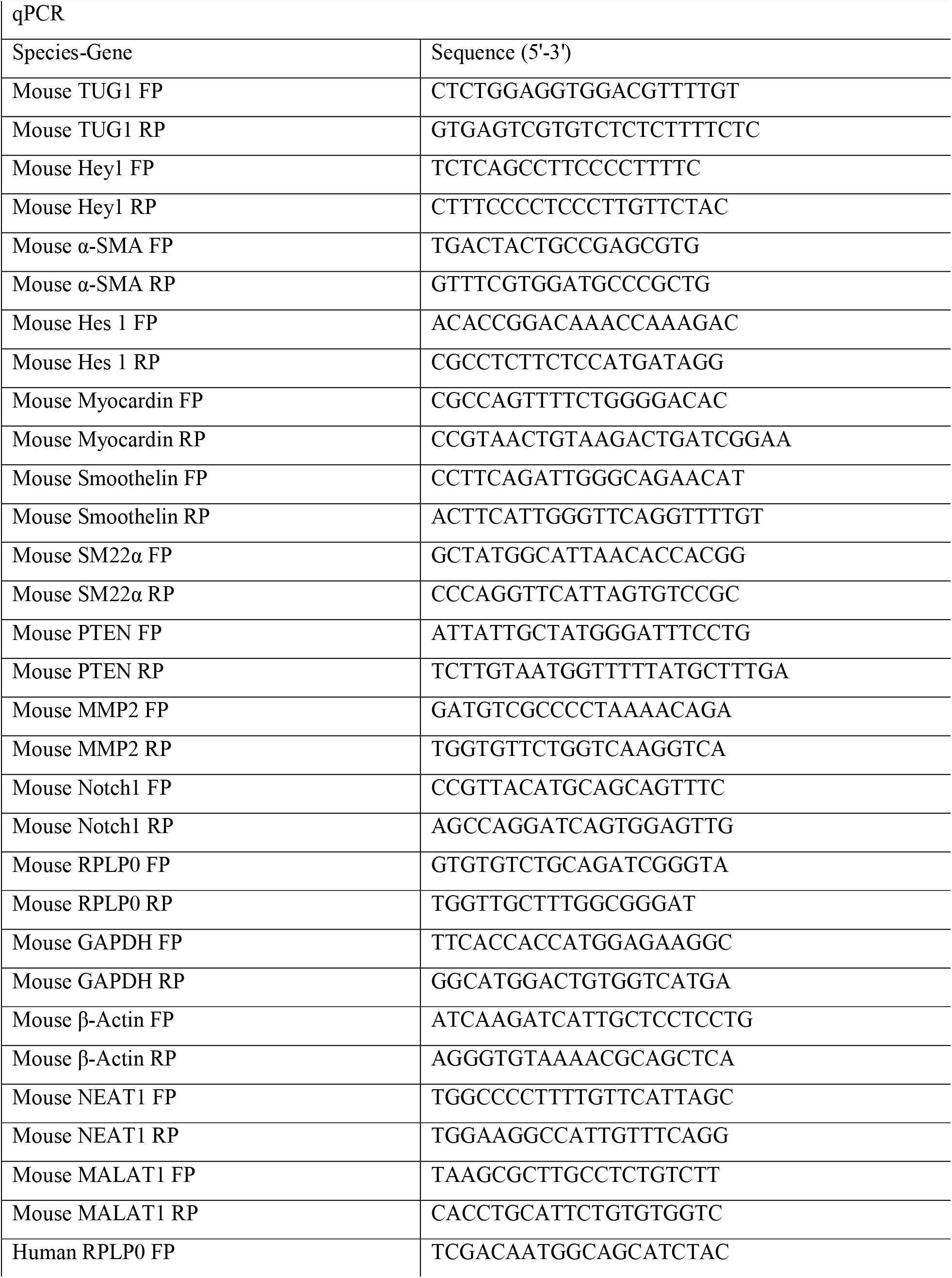
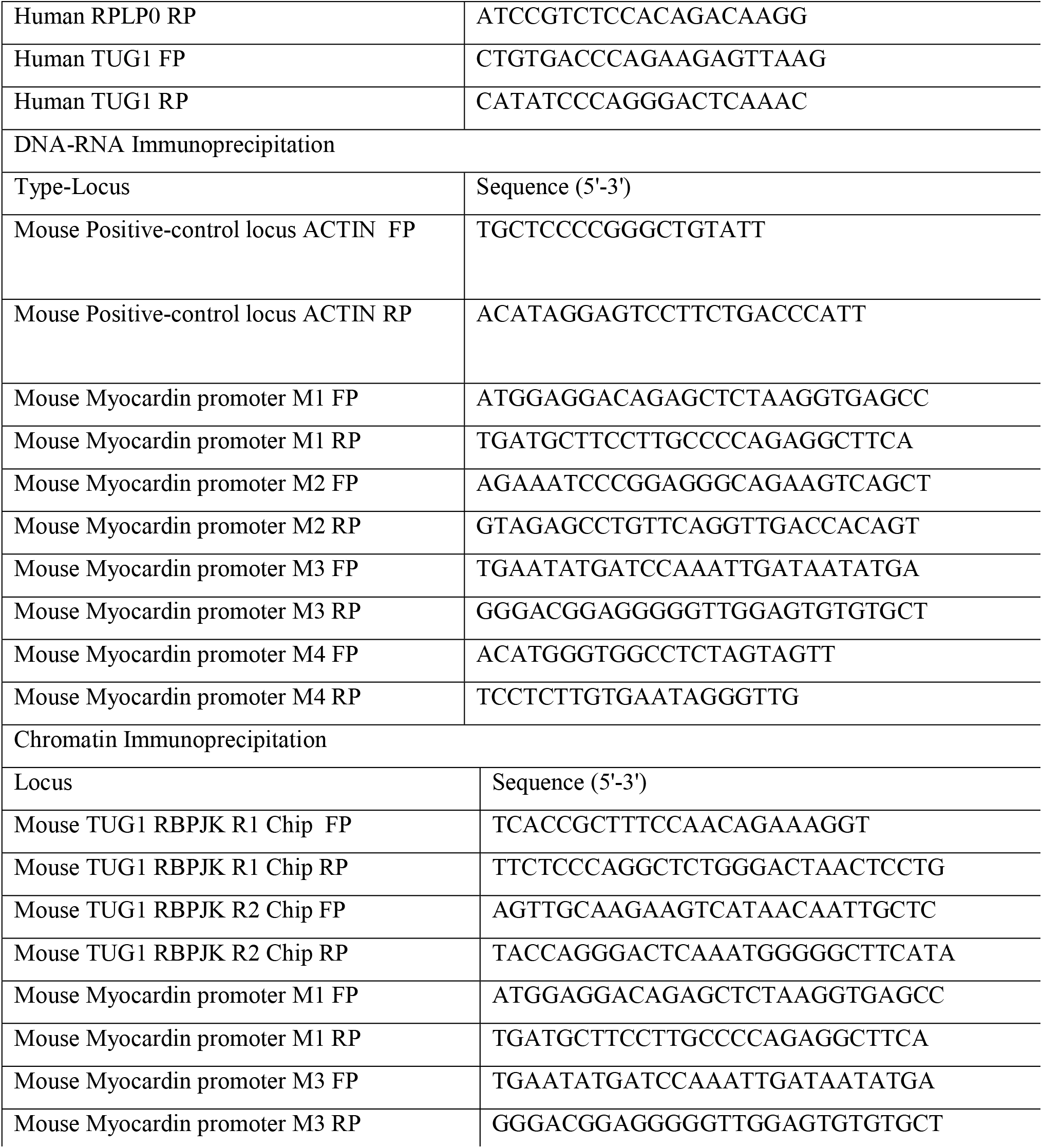
List of primers.

### Chromatin Immunoprecipitation (ChIP), RNA Immunoprecipitation (RIP) and DNA-RNA Immunoprecipitation (DRIP)

ChIP experiments were performed in mouse aortic SMCs (MOVAS) as per established protocol (Gade and Kalvakolanu, 2012) with few modifications, using anti-Notch1-ICD (Abcam, Cat no ab8925) or anti-KLF4 antibody (Novus Biologics Cat no AF3158). Upon immunoprecipitation, DNA was isolated by phenol:chloroform:isoamyl alcohol extraction and qPCR was performed using primers spanning RBPJK binding sites on TUG1 promoter. For chromatin pulldown with KLF4, primers spanned Myocardin promoter. Fold enrichment was calculated relative to percent-input using previously published method (Solomon, Caldwell and Allan, 2021). Primers used are mentioned in Table 1. Antibody dilutions used are mentioned in Table 2.

**Table 2.** List of Antibodies and dilutions.

| Antibody | Make and Cat No | Purpose | Dilution |
| --- | --- | --- | --- |
| Rabbit Anti-Calponin Antibody | Abcam - ab46794 | Immunofluorescence | (1:1000) |
| Rabbit Anti-Elastin antibody | Abcam - ab21610 | Immunofluorescence | (1:200) |
| FITC Donkey anti-Goat | Jackson ImmunoResearch - 705-093-003 | Immunofluorescence | (1:500) |
| FITC Goat anti-Rabbit | Jackson ImmunoResearch - 111-095-003 | Immunofluorescence | (1:1000) |
| Rabbit Anti-α-smooth muscle actin | Abcam - ab5694 | Immunofluorescence | (1:1000) |
| Goat Anti-SM22α | Novus Biologics- NB600-507 | Immunofluorescence | (1:500) |
| Rabbit IgG, polyclonal Antibody | Abcam - ab171870 | Chromatin Immunoprecipitation | 2μg per 10μg Chromatin |
| Rabbit Anti-activated Notch1 Polyclonal Antibody | Abcam - ab8925 | Chromatin Immunoprecipitation | 2μg per 10μg Chromatin |
| Goat Anti-KLF4 antibody | Novus Biologics- AF3158 | Chromatin Immunoprecipitation | 1μg per 10μg Chromatin |
| Goat IgG Isotype Control | Novus Biologics - AB-108-C | Chromatin Immunoprecipitation | 1μg per 10μg Chromatin |
| Anti-DNA-RNA Hybrid Antibody, clone S9.6 | Merck - MABE1095 | Chromatin Immunoprecipitation | 3.5μg per 15μg Chromatin |

**Table 3.**
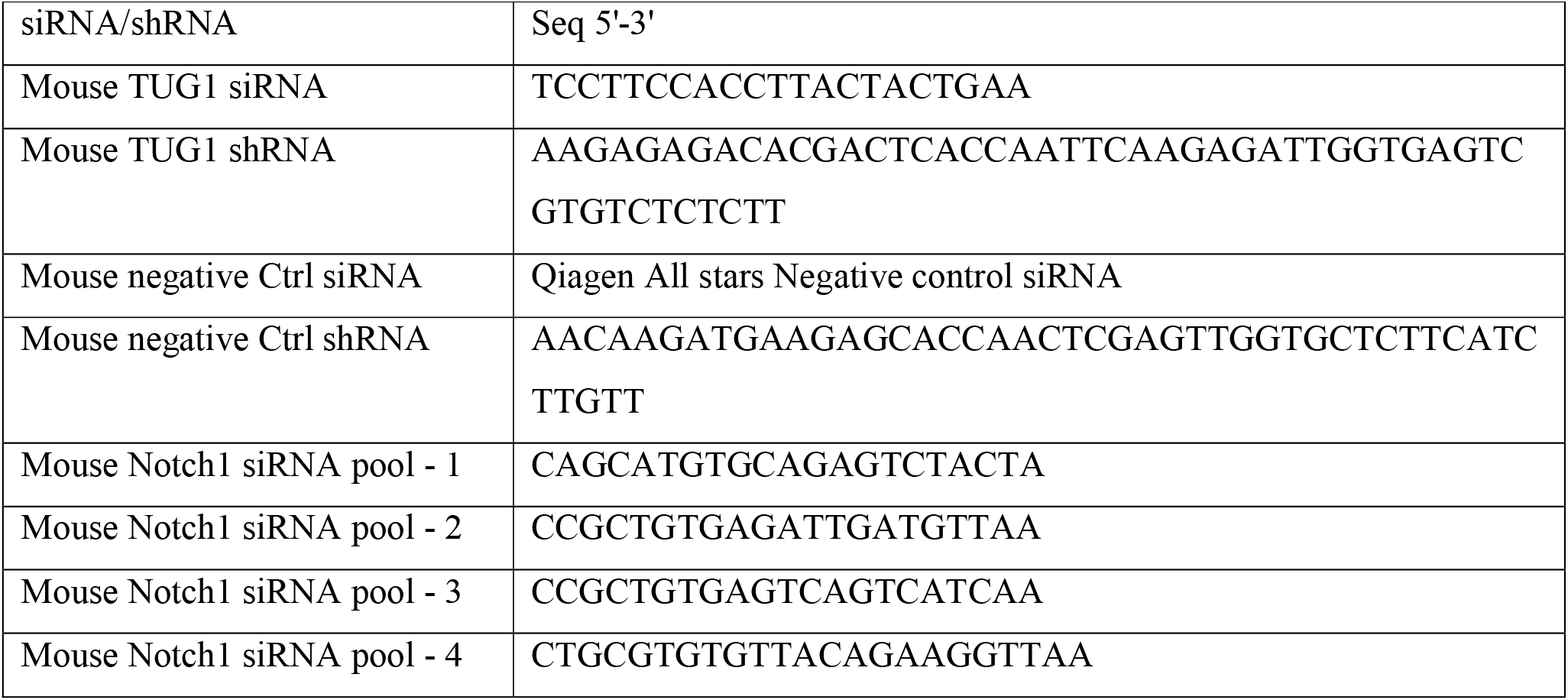
Sequence of si-RNAs and shRNA.

RNA immunoprecipitation was performed using anti-KLF4 antibody (Novus Biologics, Cat No – AF3158) in mouse aortic SMCs (MOVAS) as described earlier (Bittencourt and Auboeuf, 2012). After elution, RNA was isolated using Trizol reagent and RNA was resuspended in nuclease free water. To get rid of DNA contaminants, RNA was digested with DNase I (NEB) and cDNA was synthesized. qPCR was performed using primers to detect TUG1 RNA. Fold enrichment relative to percent-nuclear input was calculated using previously described (Solomon, Caldwell and Allan, 2021). MALAT1 and β-Actin were used as control RNA. Primers used are mentioned in Table 1. Antibody dilutions used are mentioned in Table 2.

DNA-RNA Immunoprecipitation was performed using Anti-DNA-RNA Hybrid Antibody S9.6 (Merck Cat No MABE1095) as per Sanz and Chédin, 2019. Briefly, DNA isolated from lysed mouse aortic SMCs was digested with enzyme cocktail containing HindIII – XbaI – EcoRI (NEB). Following digestion, DNA was isolated by phenol:chloroform:isoamyl alcohol extraction. Quality of digestion was indicated by fragment size of 500 bp - 1000 bp verified on agarose gel. For RNAse-H control, fragmented DNA was treated with RNAse-H (NEB) for 4-6 h at 37 °C. Prior to addition of antibody, 10 % sample was separated as Input. Samples were then incubated with antibody overnight on a rotospin at 4 °C and subsequently antibody bound DNA-RNA hybrids were captured on Protein A-Protein G beads. DNA-RNA hybrids were eluted and treated with Proteinase K (VWR Scientific). Eluted DNA was subsequently isolated by phenol:chloroform:isoamyl alcohol extraction and DRIP-qPCR was performed on Myocardin promoter and control locus and results were calculated as per Sanz and Chédin, 2019. Primers used are mentioned in Table 1. Antibody dilutions used are mentioned in Table 2.

### Statistics

All data were expressed as mean ± SD values and p-values were calculated by two tailed unpaired t test using Graphpad Prism 7. Significant differences are indicated as *p<0.05, **p<0.01, ***p<0.001, ****p<0.0001. All cell culture experiments were performed with 3-4 biological replicates and have been performed at least thrice independently. All animal studies were performed with atleast three animals per group. Promoter luciferase experiments were performed twice independently. All immunoprecipitation studies were performed atleast two times and error bars reflect independent experiments. The number of replicates used in experiments performed is also mentioned in each figure.

## Acknowledgements

This study was supported by DBT/Wellcome Trust India Alliance (India Alliance) fellowship grant to RK (fellowship No. IA/I/16/1/502357). RAB received fellowship from Council of Scientific and Industrial Research (CSIR). We thank Dr. Avinash Raj (CSIR-Centre for Cellular and Molecular Biology) for assistance with histology, Dr M Jerald Mahesh Kumar, N Sai Ram and S Prashanth (CSIR-Centre for Cellular and Molecular Biology) for assistance and expertise with animal studies. Core facilities at CSIR-Centre for Cellular and Molecular Biology are acknowledged for their technical assistance.

## Contributions

Conceptualization: R.K and R.A.B, Methodology: R.K and R.A.B, Investigation: R.A.B, Data curation: R.A.B, Resources: R.A.B, Writing and editing: R.K and R.A.B, Supervision: R.K, Funding acquisition: R.K

## Supplemental Figures

**Supplementary Fig 1.**
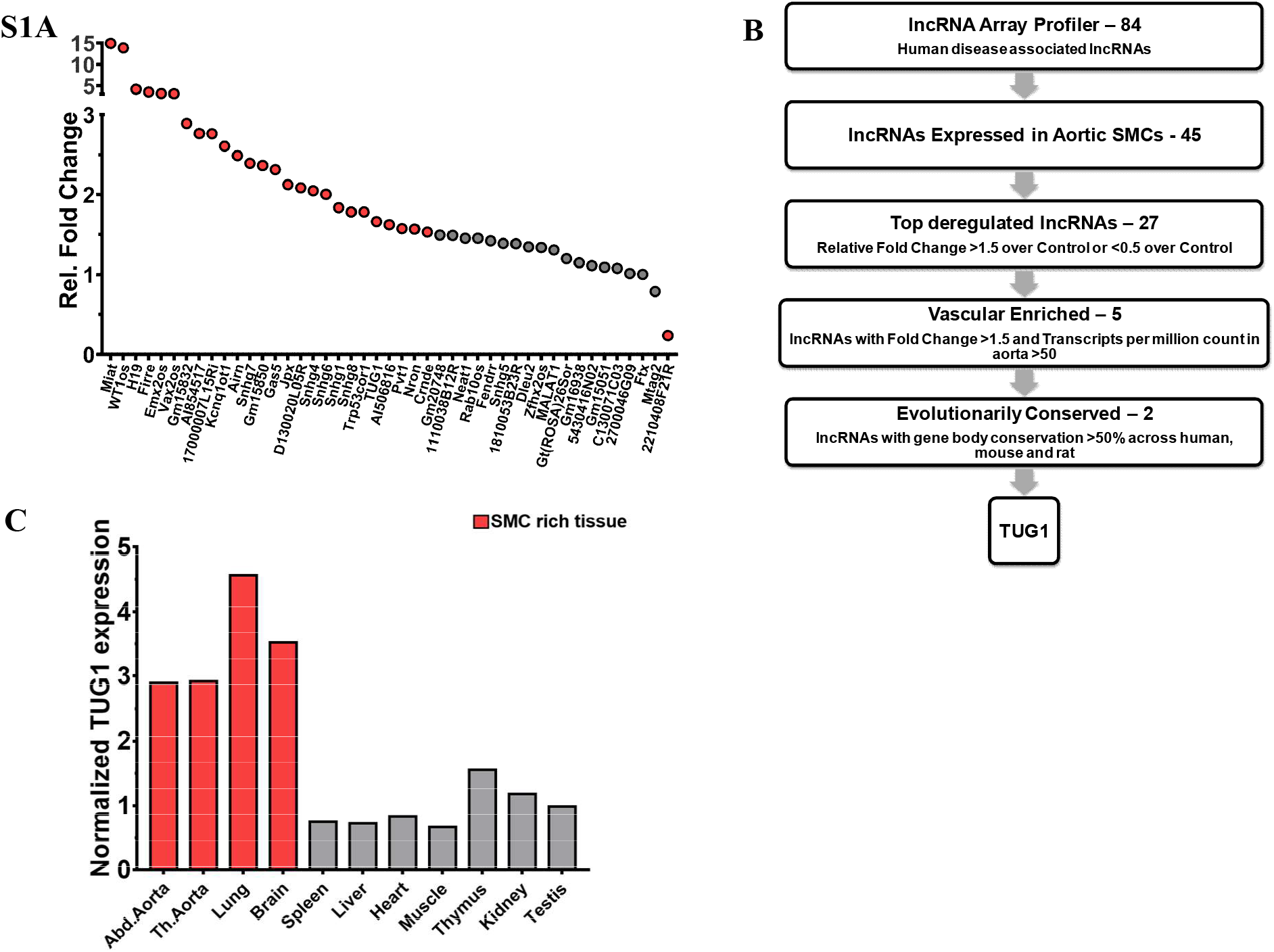
(A) LncRNA array prolifing of Ang-II induced lncRNAs in mouse aortic SMCs (MOVAS), data is shown as relative fold change over control normalized to Rns7k gene, deregulated genes are shown in red (Fold change >1.5 or Fold change <0.5). (B) Sorting criteria for candidate lncRNA expressed in lncRNA array profiler. (C) TUG1 expression analysis was performed across mouse organ library, data is shown as TUG1 expression normalized to RPLP0 of each organ, SMC rich tissues are shown in red.

**Supplementary Figure 2.**
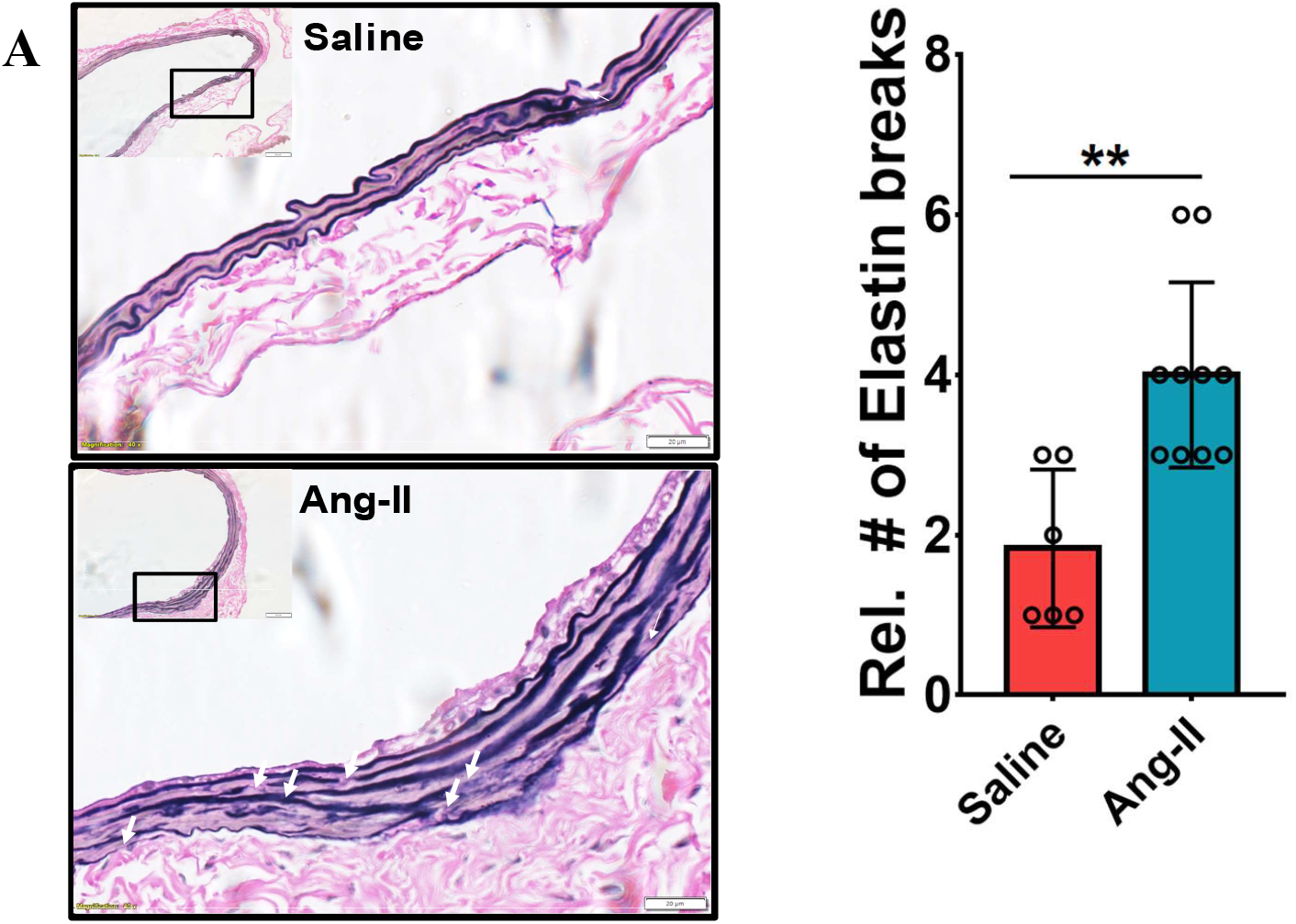
ECM degradation and histological changes during Angiotensin-II induced Abdominal Aortic Aneurysm in ApoE^-/-^ mice. (A) Representative image of cross section of verhoff van gieson stained abdominal aortas with Elastin breaks (indicated as white arrows) and quantification of elastic breaks, data is shown as relative # of breaks in elastin bands quantified between saline and aneurismal abdominal aortas, two tailed unpaired t test was used to determine statistical significance using 8 serial sections per group (right). * Denotes p<0.05 ** denotes p<0.01 **** denotes p<0.0001 unpaired t test vs Saline, Bar 20 μm.

**Supplementary Figure 3.**
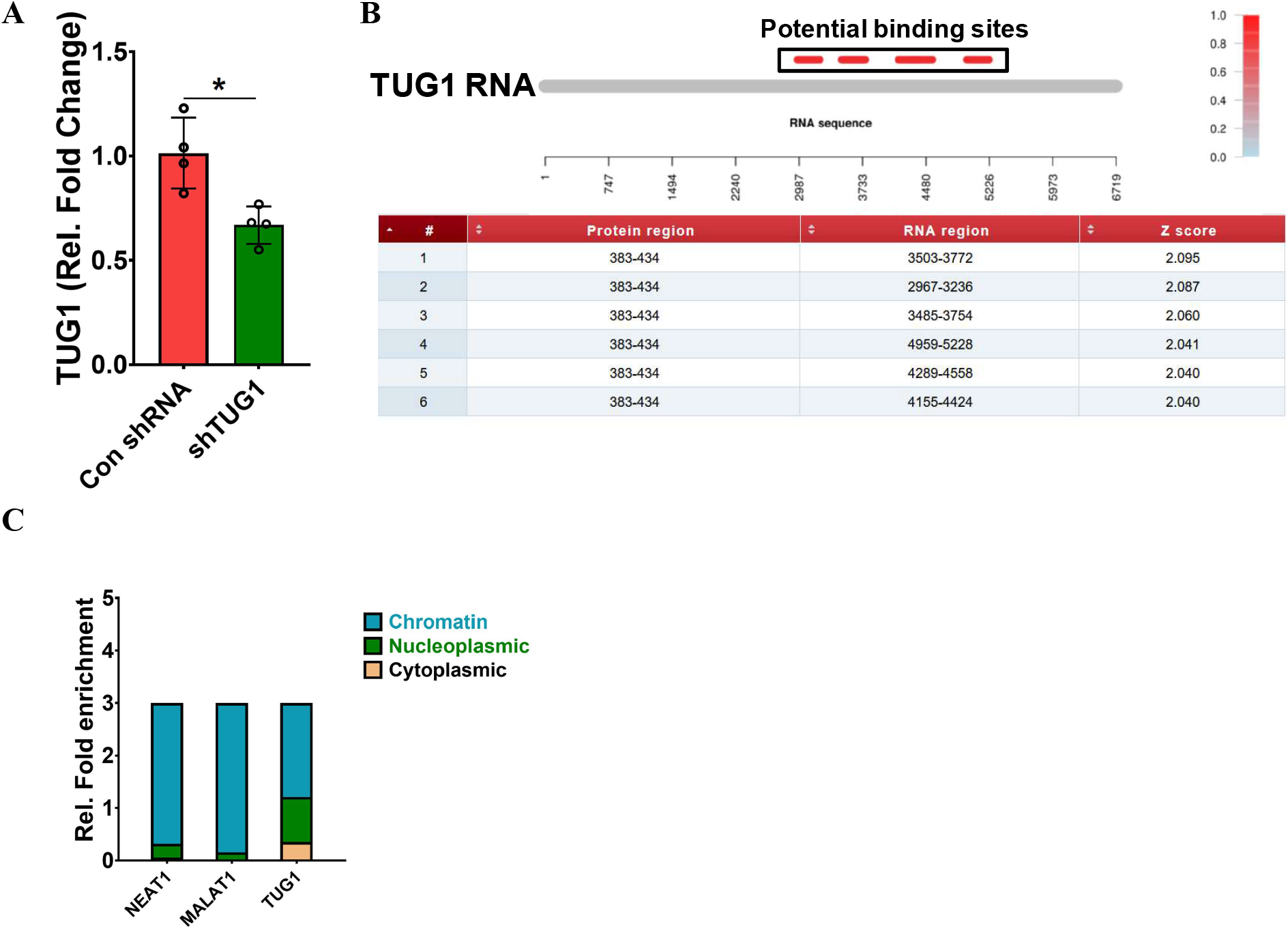
(A) shRNA mediated KD of TUG1 in mouse aortic SMCs (MOVAS), data is shown as relative fold change over control shRNA normalized to RPLP0 and depicted as mean±SD, n=4/group. (B) catRAPID interaction propensity of TUG1 and KLF4 with interaction score and amino acid-nucleotide residue numbers. Red bars above TUG1 transcript denote potential binding regions. (C) Sub-cellular Fractionation of TUG1 in SMCs. NEAT1 and MALAT1were used as positive controls for nuclear fraction. * Denotes p<0.05, unpaired t test vs Con-shRNA.

**Supplementary Table 1:** Vascular enriched long non coding RNAs (lncRNAs) are shown as lncRNAs with Transcripts per million (TPM) > 50 in human aorta derived from GTEx database and fold change >1.5 in Ang-II treated mouse aortic SMCs.

| lncRNA | TPM in human aorta |
| --- | --- |
| Snhg6 | 196.3 |
| GAS5 | 140.2 |
| Snhg8 | 106.5 |
| TUG1 | 80.63 |
| Snhg1 | 50.63 |

